# Comparison of human and mouse fetal intestinal tissues reveals differential maturation timelines

**DOI:** 10.1101/2020.06.18.157818

**Authors:** A.A. Lim, R.R. Nadkarni, B.C. Courteau, J.S. Draper

## Abstract

Maturation of the intestinal epithelium is a necessary step for development of a fully functioning gastrointestinal tract. Studies of rodent gastrointestinal development and maturation have long been used to guide understanding of human intestinal maturation, in part because accessing human gestational stage intestinal tissues to perform equivalent human studies can be difficult. Notable differences have already been described in the timing of key stages in intestinal development between rodents and humans, but the conservation of intestinal maturation events between the two species is poorly understood. We hypothesized that species-related differences in intestinal development would alter the timing of key maturation events between human and mouse. We tested our hypothesis by performing a detailed comparison of hallmarks of intestinal maturation in human and mouse gestational intestine, including markers that describe the emergence of intestinal cell types, functionality and structural integrity. Our study demonstrates clear timing differences between maturation stages in mouse and human, with the majority of human maturation hallmarks acquired post-partum, in contrast to their gestational emergence in mouse. Our work suggests caution when translating murine intestinal maturation observations to the human, and provides a maturation road map that will be helpful to those seeking to produce mature intestine from *in vitro* stem cell sources.

## Introduction

Intestinal maturation plays a critical role in the normal development and function of the native intestine. Failure of the intestine to properly mature is implicated in the pathogenesis of neonatal intestinal diseases like necrotizing enterocolitis (NEC) and early-onset inflammatory bowel disease (IBD), in which bacteria invade the intestine, resulting in inflammation and potential destruction of the wall of the intestine (Abraham and Cho, 2009; Neu and Walker, 2011). In fact, NEC is the most common disease among premature infants, affecting about 7% of cases, and with a 25% mortality rate (Rich and Dolgin, 2017). An improved comprehension of human intestinal maturation is important for identifying the factors that contribute to the development of a healthy functional intestine. Understanding the hallmarks of human intestinal maturation will also assist in the production of intestinal tissues from pluripotent stem cell sources, which are typically characterized by an immature phenotype.

Intestinal maturation can be defined as functional changes that occur beyond morphogenesis and differentiation into intestinal cell types, and is marked by the emergence or loss of key protein markers, improvement of structural integrity, and host defense. For example, when enterocytes first appear in the intestine, they express villin, but functionality coincides with the production of glucosidase enzymes such as sucrase isomaltase, trehalase, and lactase (Finkbeiner et al., 2015). Similarly, emerging Paneth cells secrete lysozyme, but are only considered mature when they secrete α-defensin peptides for host defense (Finkbeiner et al., 2015; Mallow et al., 1996). The expression of tight junction protein Claudin 3 in the intestinal epithelium is an indicator of improved epithelial barrier integrity and reduced intestinal permeability (Günzel and Yu, 2013; Lu et al., 2013; Milatz et al., 2010; Patel et al., 2012). Failure or untimely emergence of such changes can lead to impaired intestinal function.

Much of the current understanding of human intestinal development and maturation has been enriched by and extrapolated from studies in the mouse. However, the timing of key events that shape intestinal development and maturation in human is now understood to deviate from that observed in the mouse. Not only is the length of the gestational period itself vastly different between the two species, lasting only 21 days in mouse but taking around 36 weeks in human, but the relationship between equivalent gestational ages is not linear (Otis and Brent, 1954). In mouse, intestinal morphogenesis is completed several weeks after birth, whereas human morphogenesis is completed several weeks prior to birth **(Figure 1)** (McCracken and Lorenz, 2001). These developmental differences are already understood to be underpinned by the differential emergence of cell types and indicators of intestinal maturation when contrasting mouse and human. For example, intestinal crypts and Paneth cells emerge around postnatal day 14 in the murine intestine, equivalent to around week 20 of gestation in the human intestine when they emerge in the latter. However, similarities between mouse and human in the timing of important intestinal maturational milestones such as the tightening of the epithelial barrier, which occurs several weeks after birth in mice (Patel et al., 2012), still require clarification. Here, we sought to provide insights into the timing of critical hallmarks of intestinal development and maturation in human tissue. We assessed the expression of protein markers of maturation using tissue sections of gestational human intestine and comparing with that of mouse. We found that, while majority of maturation markers emerge several weeks after birth in the murine intestine, the same hallmarks were visible prior to birth in human. Our study provides valuable insights into intestinal development and maturation.

**Figure 1.**
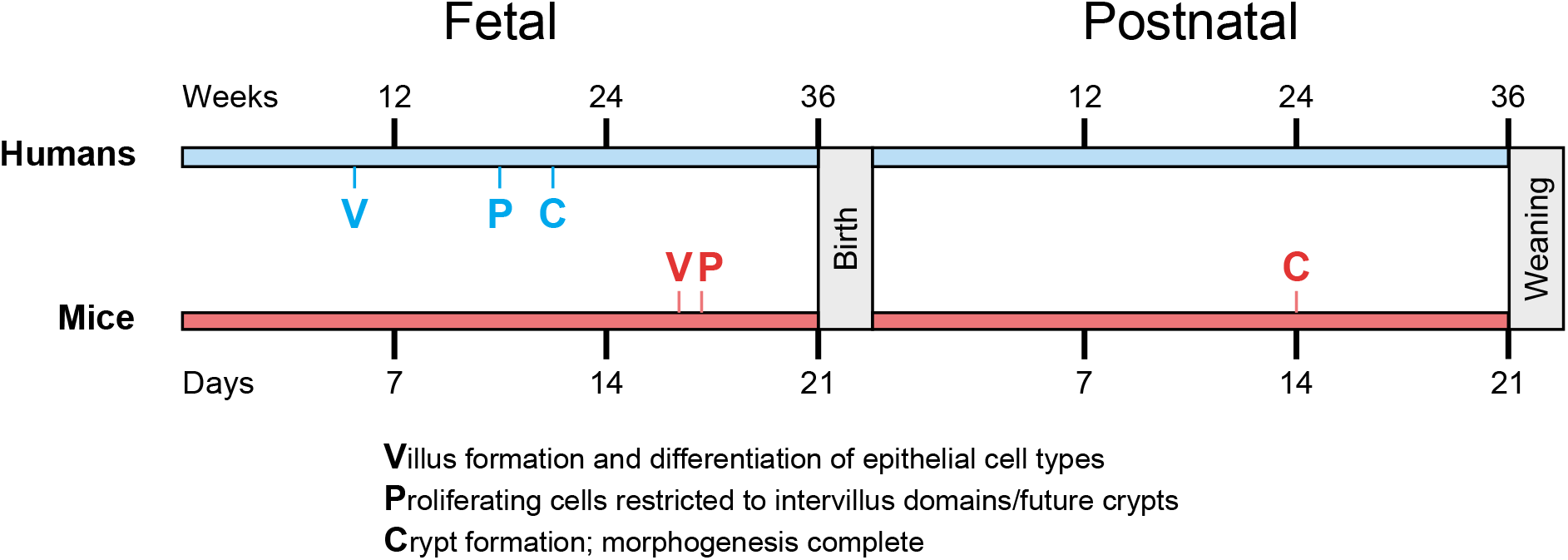
A comparison of the timeline between human and murine intestinal epithelial development from conception to weaning. Morphogenesis of the intestinal epithelium in humans is complete several months before birth around the end of the second trimester, whereas in mice it is complete around two weeks after birth (McCracken and Lorenz, 2001).

## Results

To investigate human intestinal organogenesis, samples from weeks 13, 18, 24 and 32 of gestation, corresponding to the end of the first trimester, middle and end of the second trimester, and middle of the third trimester, respectively, as well as adult intestinal tissue, were obtained under REB approval. Comparative tissue sections of murine intestine were sourced at E17.5, P7, P14, P21 and P60, which range from post-villus formation to post-weaning and mature adult intestine. CDX2 protein expression (Silberg et al., 2000; Zorn and Wells, 2009) **(Figure S1)** and haematoxylin and eosin (H&E) staining **(Figure S2)** were used to verify intestinal identity for all samples in this study. H&E staining revealed differences in human intestinal epithelial structure with age, such as the absence of crypt-resembling domains at weeks 13 and 18 but their presence from week 24 onwards **(Figure S2)**, supporting the observation that crypts emerge around week 20 in the human intestine (Chin et al., 2017). Alcian blue staining demonstrated that mucin-secreting goblet cells were present in the epithelium as early as week 13 in humans and E17.5 in mice **(Figure S2)**.

Human and mouse tissues were next compared for key proteins that mark cell type emergence, functionality, structural integrity, and protective role, and that together constitute intestinal maturation (Chin et al., 2017; Finkbeiner et al., 2015). A description of the role of each marker assessed in this study is summarized in **Table 1**.

**Table 1.** A summary of molecular indicators of intestinal maturation in this study.

| **Protein marker** | **Type of protein** | **Description of identity in maturation** | **References** |
| --- | --- | --- | --- |
| Chromogranin A (CHGA) | Neuroendocrine secreted protein | Secreted by enteroendocrine cells | Gunawardene et al., 2011; Noah et al., 2011 |
| Claudin 3 (CLDN3) | Tight-junction protein | Indication of improved epithelial barrier integrity and reduced intestinal permeability | Günzel and Yu, 2013; Lu et al., 2013; Milatz et al., 2010; Patel et al., 2012 |
| Defensin, alpha 1 (DEFA1) in mice, or Defensin, alpha 5 (DEFA5) in humans | Host defense peptide | Secreted by functional Paneth cells, and involved in host defense | Ayabe et al., 2000; Ayabe et al., 2002; Finkbeiner et al., 2015 |
| Olfactomedin 4 (OLFM4) | Anti-apoptotic factor; extracellular matrix glycoprotein | Mature stem cell marker; becomes expressed in crypt base columnar cells in mature crypts | van der Flier et al., 2009; Finkbeiner et al., 2015 |
| Sucrase isomaltase (SI) | Glucosidase enzyme | Produced by mature enterocytes | Finkbeiner et al., 2015 |
| Ulex Europaeus agglutinin 1 (UEA1) | Glycoprotein-binding lectin | Binds to fucosylated glycoproteins of enterocytes, goblet cells, and Paneth cells; has protective role in gut infection | Pickard and Chervonsky, 2015 |

### Sucrase isomaltase and chromogranin A, markers of brush border and enteroendocrine maturation respectively, appear during early gestation in humans

Villus morphogenesis begins at E15 in mice, and is one of the first signs of differentiation in the developing intestine (Chin et al., 2017; Noah et al., 2011). Absorptive enterocytes are one of the intestinal epithelial cell types that emerge during this period, at around E16.5. Enterocytes comprise more than 80% of the intestinal epithelium and are responsible for absorbing nutrients from the lumen. Due to the presence of microvilli on their apical surface, enterocytes constitute the brush border of the intestine (Chin et al., 2017). Villin (VIL1) is an actin-binding protein that is expressed in the brush border throughout development and in the adult. However, a fully functional brush border is distinguished by the production of glucosidase enzymes, such as sucrase isomaltase (SI), whose expression is low in the fetal intestine but peaks in the adult intestine in humans (Finkbeiner et al., 2015). Therefore, SI expression is thought to be an indicator of brush border maturation, but it is unclear when it emerges at the protein level in the intestine. SI was visible in mice only at P21 and not earlier **(Figure 2)**, even though villus formation and differentiation is complete by E17. In contrast, SI was clearly expressed in all tested gestational ages in human including week 13 **(Figure 2)**, demonstrating that SI protein expression in the human intestine begins as early as the end of the first trimester, challenging the notion that it emerges later as a marker of brush border maturation (Finkbeiner et al., 2015).

**Figure 2.**
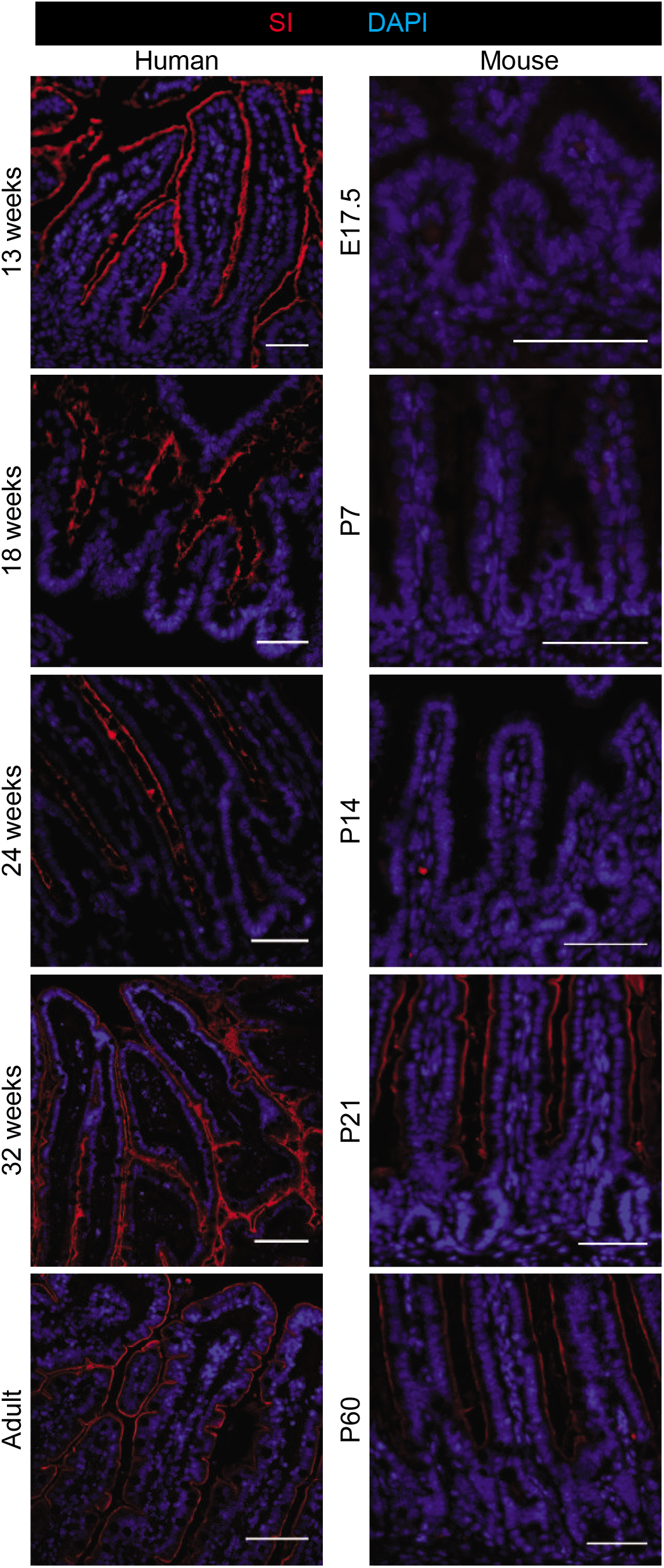
Fluorescent IHC staining of gestational and postnatal small intestine in humans versus mice for SI. Scale bars, 50 μm.

Intestinal enteroendocrine cells (EECs) are specialized cells with endocrine function that produce hormones in response to various stimuli and release them into the bloodstream or to the enteric nervous system. In adults, EECs comprise ∼1% of the intestinal epithelium and are located sporadically throughout (Gunawardene et al., 2011; Noah et al., 2011). EECs emerge at around E16.5 in mice shortly after villus morphogenesis begins (Chin et al., 2017), but it is unclear how early EECs emerge in the human intestine. EECs are considered a putative indicator of maturation due to the difficulty in generating them *in vitro* in human pluripotent stem cell (hPSC)-derived intestinal tissues which tend to be fetal-like (Fordham et al., 2013; Forster et al., 2014; Nadkarni et al., 2017). The expression of Chromogranin A (CHGA), a neuroendocrine secretory protein, marks EECs. CHGA-expressing cells were present in all human gestational ages tested, including week 13 in the developing epithelium **(Figure S3), i**ndicating that intestinal EECs emerge as early as the end of the first trimester in humans.

### The intestinal stem cell marker OLFM4 is expressed in human crypt base columnar cells before birth

Intestinal crypt formation in mice begins shortly after birth and is complete by P14, whereas human crypt formation occurs before birth, at around week 20 of gestation (Chin et al., 2017). Lgr5-expressing intestinal stem cells reside in the crypts and are capable of differentiating into other intestinal epithelial cell types in response to wear and tear and injury (Noah et al., 2011; Umar, 2010). Olfactomedin 4 (OLFM4) is a robust marker of Lgr5-type crypt base columnar cells in the adult intestine, and is an indicator of crypt maturation (Finkbeiner et al., 2015; van der Flier et al., 2009). We hypothesized that emergence of OLFM4 in the crypts coincides with crypt morphogenesis in both mice and humans. In mice, OLFM4 was expressed in crypt-like domains from P7 onwards but not earlier **(Figure 3A)**. Human OLFM4 expression was detected in the crypts by week 24 of gestation, and was clearly localized to crypt base columnar cells at week 32 and in the adult **(Figure 3A)**. OLFM4 staining was not visible at weeks 13 and 18 **(Figure 3A)**, consistent with the absence of crypts prior to week 20. Therefore, the emergence of OLFM4 expression coincides with crypt formation, and the localization of OLFM4 in the base of the crypt begins around the end of the second trimester.

**Figure 3.**
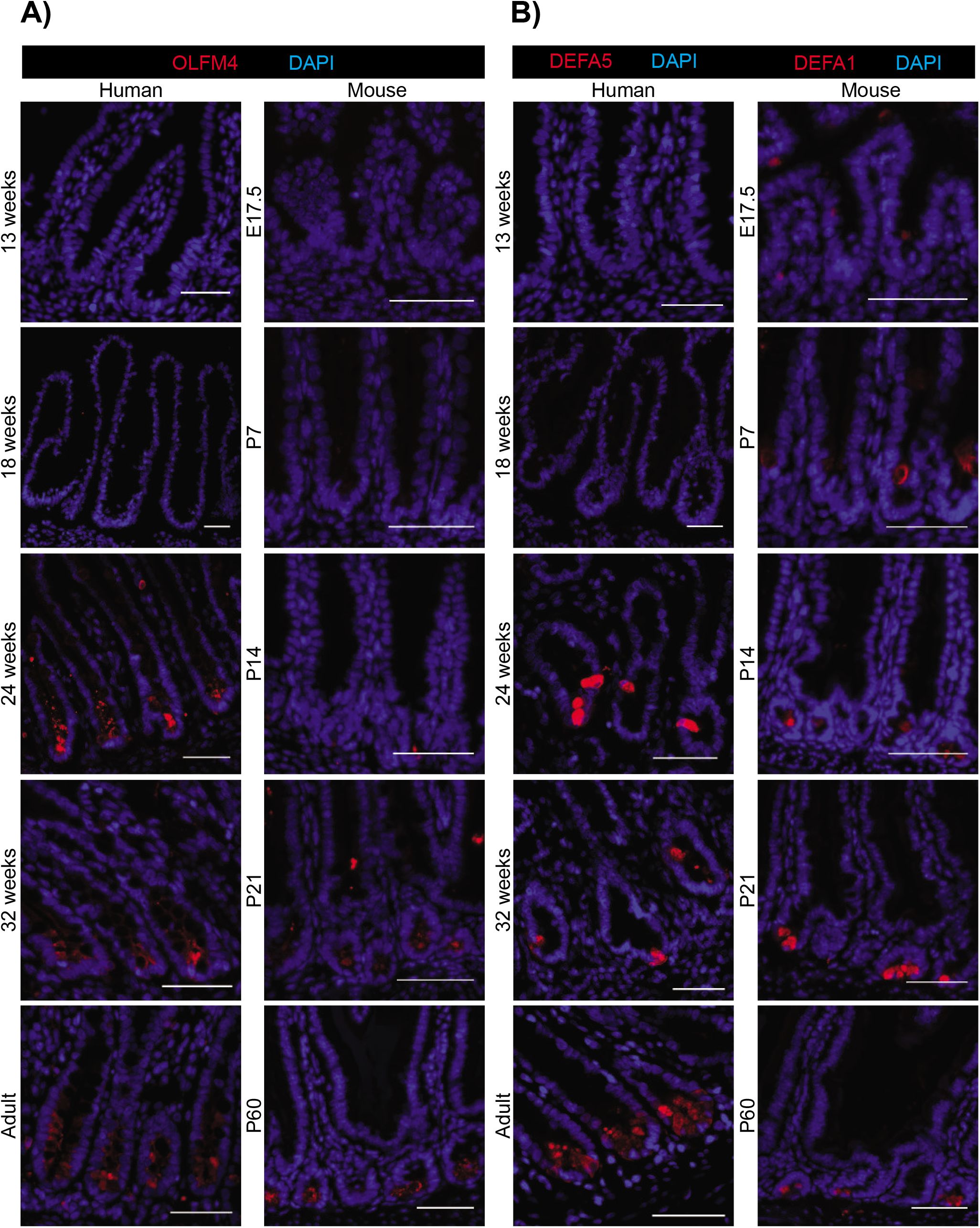
Fluorescent IHC staining of gestational and postnatal small intestine in humans versus mice for **A)** OLFM4 and **B)** DEFA5(h)/DEFA1(m). Scale bars, 50 μm.

### Expression of α-defensin 5, a marker of Paneth cell maturation, emerges prior to birth in the human intestine

Paneth cell differentiation coincides with intestinal crypt emergence (Calvert and Pothier, 1990; Chin et al., 2017; Kim et al., 2012), which occurs at P14 in mice and week 20 of gestation in humans (Mallow et al., 1996; Moxey and Trier, 1978). While lysozyme production marks the presence of Paneth cells in the intestine, the production of α-defensin peptides for defense against pathogens is an indicator of Paneth cell maturation and functionality (Ayabe et al., 2000; Finkbeiner et al., 2015). Since Paneth cells appear around mid-gestation in the human intestine, we hypothesized that Paneth cell maturation also occurs around this time. We tested this hypothesis by staining for α-defensin 5 (DEFA5 or HD5) in humans. Due to a lack of suitable commercially available antibodies with reactivity against murine DEFA5, murine intestinal sections were stained for α-defensin 1 (DEFA1), an α-defensin peptide with similar functionality (Ayabe et al., 2000; Mallow et al., 1996). DEFA1 was expressed in the developing mouse crypts only from P7 onwards and was fully localized to the crypts by P14 **(Figure 3B)**. In humans, DEFA5-expressing cells were present in the crypts at weeks 24 and 32 of gestation, appearing to be progressively restricted to the crypt base over time **(Figure 3B)** (Bjerknes and Cheng, 1981; Kim et al., 2012). DEFA5 staining was not visible at weeks 13 and 18 **(Figure 3B)**, consistent with the absence of crypts and Paneth cells before week 20. These observations suggest that Paneth cell maturation occurs before birth soon after Paneth cell specification in the human intestine.

### Proliferating cells become confined to intervillus domains as early as week 13 in the human intestine

Prior to and during villus formation (at E15 in mice), proliferating cells are scattered throughout the developing epithelium. After villus formation, proliferating cells progressively decline in the villi, and by E17, they become confined to the intervillus domains, which later become crypts at the base of the villi for the remainder of development. The crypts emerge from the intervillus domains around P14 in mice and week 20 of gestation in humans, containing all stem and proliferating cells of the mature intestinal epithelium (Chin et al., 2017; Noah et al., 2011). Based on our above data in humans which shows that indicators of villus differentiation are present as early as week 13, we hypothesized that proliferating cells marked by Ki67 would already be confined to intervillus domains at this stage. We found that Ki67-expressing cells were largely confined to intervillus domains at week 13, completely confined at week 18, and later in crypts from week 24 onwards **(Figure S4)**. These results are consistent with the trend observed in the murine intestine, although with a shifted timeline.

### Fucosylation, marked by UEA1 reactivity, appears to begin before birth in the human and murine intestine, and becomes spatially localized to epithelial cell types during morphogenesis

Fucosylation, a type of glycosylation in which fucose units are added to glycoproteins and glycolipids, is thought to serve a protective role in both intestinal and systemic infection and inflammation through suppressing the virulence of harmful pathogens (Pickard and Chervonsky, 2015; Pickard et al., 2014). In the murine intestine, fucosylation is localized to glycoproteins produced by Paneth cells in the crypts, and goblet cells and enterocytes in the villi (Bry et al., 1996; Debray et al., 1981). Fucosylation is marked by increased expression and activity of fucosyltransferase enzymes, and reactivity to the lectin Ulex europaeus agglutinin 1 (UEA1) (Pickard and Chervonsky, 2015). Fucosylation of brush border and mucinous glycoproteins in the murine intestine is thought to occur primarily after birth and post-weaning, coinciding with abundant enteric bacteria and polyamines within the luminal space (Biol-N’Garagba et al., 2002; Bry et al., 1996; Pickard and Chervonsky, 2015). The timing of appearance of fucosylation in the human intestine is poorly understood, thus human and murine samples were contrasted for reactivity to fluorochrome-conjugated UEA1. In accordance with previous studies, UEA1 staining was seen in the postnatal murine brush border at P7 and in Paneth cells after P14, but reactivity was seen in goblet cells at all tested stages including E17.5 **(Figure 4A)** (Bry et al., 1996; Debray et al., 1981). In adult human intestinal tissue, UEA1 reactivity was also clearly observed in Paneth cells, goblet cells and the brush border, but was only localized to Paneth and goblet cells at weeks 24 and 32 of gestation **(Figure 4A)**. At week 18, UEA1 reactivity was seen in the lumen in patches resembling secreted mucins, and also at week 13 but without localization in any obvious morphological structures **(Figure 4A**). This indicates that fucosylation in the human and murine intestine begins during gestation and becomes localized to differentiated epithelial cell types as development progresses.

**Figure 4.**
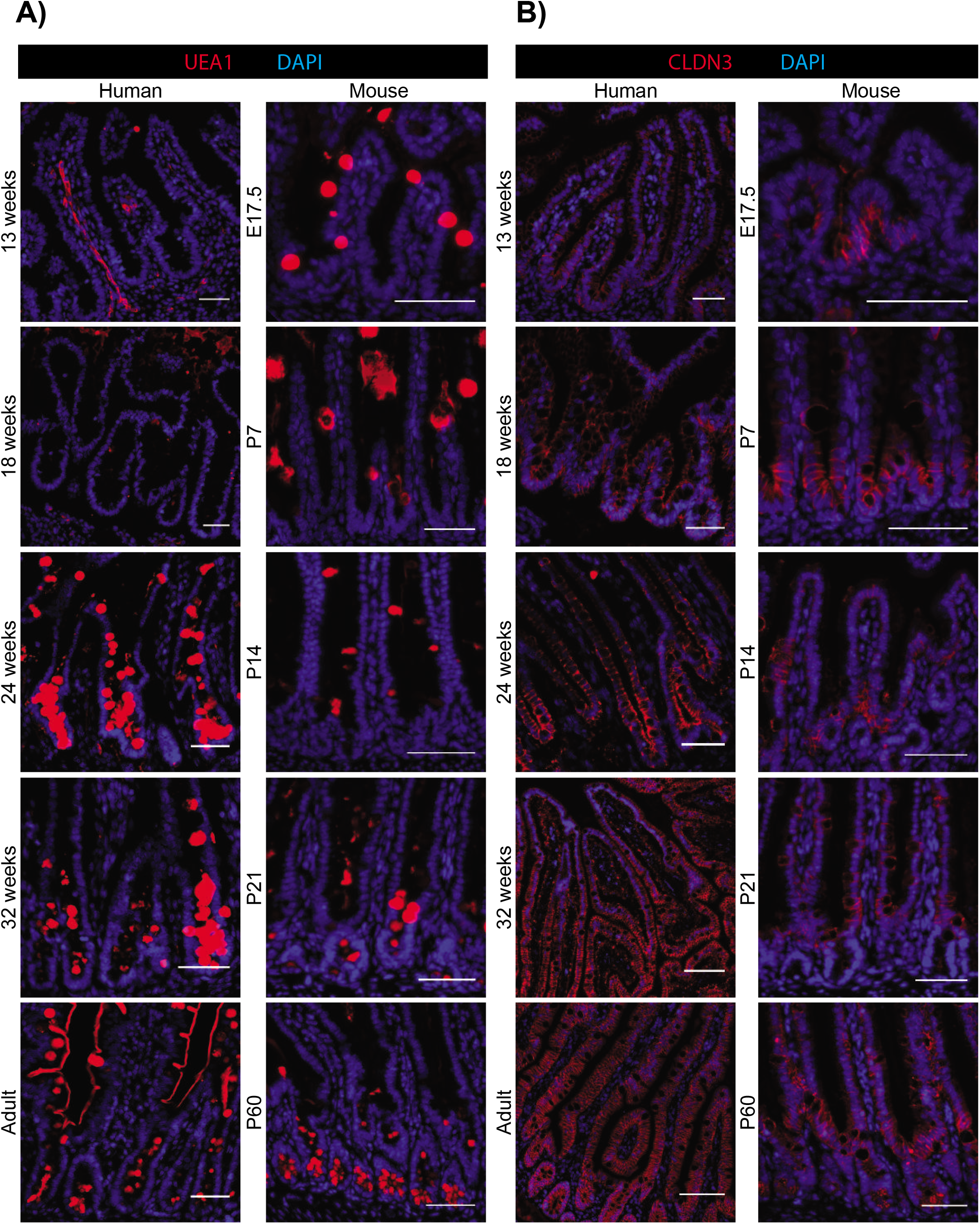
Fluorescent IHC staining of gestational and postnatal small intestine in humans versus mice for **A)** lectin UEA1 and **B)** CLDN3. Scale bars, 50 μm.

### Claudin 3, a marker of epithelial barrier maturation, is expressed before birth in the human and murine intestine

In premature infants suffering from NEC or IBD, the immature intestine is vulnerable to invading bacteria that are capable of destroying the intestinal wall. Epithelial barrier integrity is important for reducing systemic invasion of gut luminal pathogens or toxins, and the use of breast milk and probiotics can contribute to proper barrier maturation and prevention of disease (Alfaleh et al., 2011; Clayburgh et al., 2004; Grave et al., 2007; Guandalini, 2010; Klement et al., 2004; Sartor, 2008; Sisk et al., 2007). The tight junction protein Claudin 3 (CLDN3) is a marker of improved epithelial barrier integrity (Günzel and Yu, 2013; Lu et al., 2013; Milatz et al., 2010; Patel et al., 2012). In the murine intestine, CLDN3 is thought to be absent at the protein level during gestation, and becomes robustly expressed 2-3 weeks after birth when morphogenesis is complete (Patel et al., 2012). Colonization of the postnatal murine intestine with the probiotic bacterial species *Lactobacillus rhamnosus* induces CLDN3 expression in epithelial tight junctions and reduces intestinal permeability (Patel et al., 2012). We hypothesized that CLDN3 expression also emerges after birth in the human intestine as a response to bacterial colonization. In mice, CLDN3 was sporadically expressed in the developing epithelium at E17.5, became more widespread at P7 in the intervillus domains, and only at P21 did expression resemble that of the adult **(Figure 4B)**. In contrast, CLDN3 was clearly visible, albeit weakly, throughout human intestinal epithelium as early as week 18 of gestation **(Figure 4B)**. CLDN3 expression became more robustly expressed at week 24, then by week 32 more closely resembled the expression observed in the adult. This indicates that CLDN3 protein expression in the human intestine emerges before birth as early as the beginning of the second trimester.

## Discussion

Understanding tissue maturation during development can provide important insights into the processes that can contribute to diseases in early infancy and later adulthood. In addition, it has become increasingly clear that a gap in the understanding of how tissues mature is also impacting the production of cell types, including intestinal cells, from pluripotent stem cells that faithfully recapitulate adult tissue function (Finkbeiner et al., 2015; Fordham et al., 2013; Hill et al., 2017; Nadkarni et al., 2017; Spence et al., 2011). While tracking the expression of protein markers of maturation in fixed tissue sections of human intestine from different gestational ages, we observed differences to the mouse intestine for the expression of markers that have been previously described as hallmarks of maturation (Chin et al., 2017; Finkbeiner et al., 2015; Patel et al., 2012). Although transcript levels of some markers in this study have been compared between the fetal and adult human intestine (Finkbeiner et al., 2015), assessment at the protein level and at multiple gestational ages is lacking. In addition to profiling expression across gestational time points, we focused on staining sections of fetal intestinal tissue to clarify how protein localization is connected to human intestinal maturation. However, it is important to note that access to human gestational tissues of sufficient quality, especially from the first and second trimesters, limited this study to the analysis of a small number of samples from each stage. Notwithstanding, the data we provide here yielded insights into intestinal maturation events during gestation that had not previously been reported.

The emergence of EECs, which is marked by CHGA protein expression, is a putative indicator of intestinal maturation (Fordham et al., 2013; Spence et al., 2011). Our data demonstrates that CHGA is present in the developing epithelium as early as week 13 of gestation, indicating that, similar to mouse, EECs are one of the first differentiated epithelial cell types to emerge in the human intestine (Chin et al., 2017; Noah et al., 2011). The early emergence of EECs during human intestinal development raises questions as to why EECs are typically absent in intestinal tissues differentiated from hPSCs *in vitro* (Fordham et al., 2013; Forster et al., 2014; Nadkarni et al., 2017), when other cells types that arise later in development can be found. The lack of EECs could be attributed to deficiencies in exogenous signaling that activate genes responsible for promoting EEC differentiation, such as *Neurogenin-3* (Sinagoga et al., 2018; Spence et al., 2011).

Transcripts of *SI, OLFM4*, and *DEFA5*, markers of brush border, crypt, and Paneth cell maturation respectively, have been shown to be expressed at low levels in the fetal intestine and highest in the adult intestine (Finkbeiner et al., 2015). SI, OLFM4, and DEFA5 are expressed at the protein level in the adult human intestine (Finkbeiner et al., 2015), but we found that they are also present in the fetal intestine mid-gestation onwards, suggesting that brush border and crypt maturation occur before birth. Moreover, the emergence of OLFM4 and DEFA5 expression coincident with crypt formation in mice and humans underscores the difference in the timeline of intestinal morphogenesis. **Figure 5** illustrates the timeline of key events from conception to weaning and displays the emergence of maturation markers investigated in this study.

**Figure 5.**
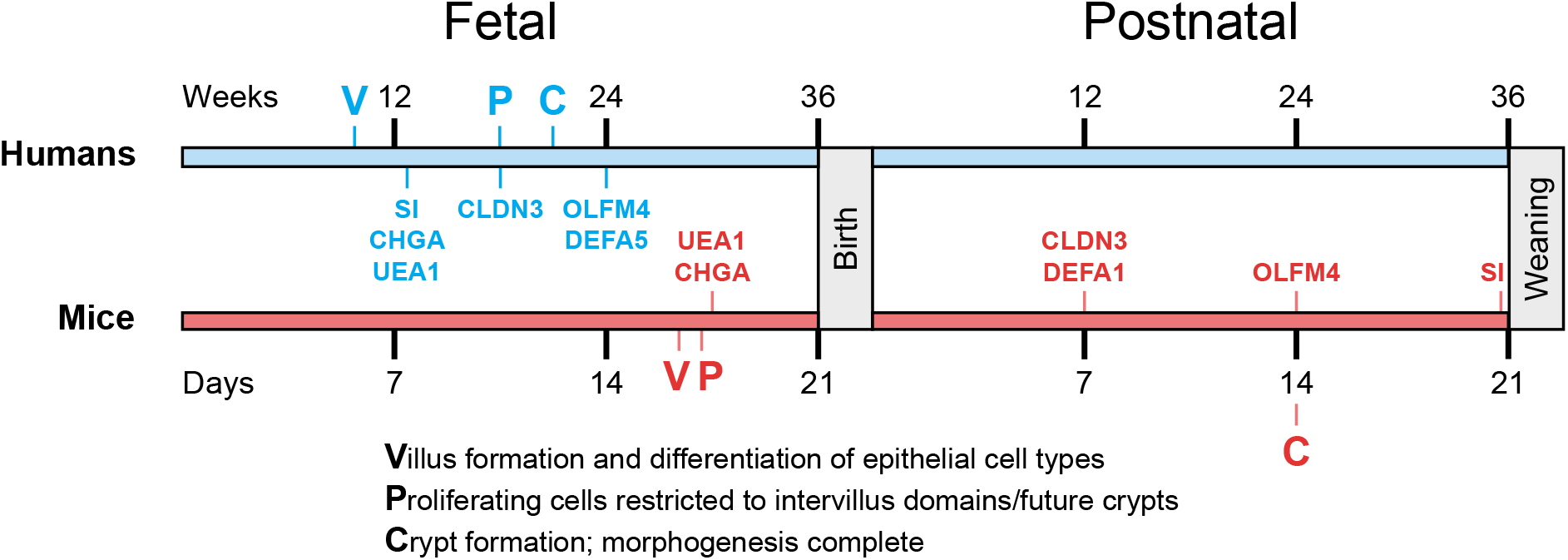
A comparison of the timeline between human and murine intestinal epithelial development from conception to weaning, displaying the emergence of maturation markers investigated in this study. Markers are placed at positions corresponding to the earliest tested human and mouse ages from which point onwards they were detected.

Integrity of the intestinal epithelial barrier, which is known to be developmentally regulated in mouse (Patel et al., 2012), is understood to be a key determinant in the transport of important molecules across the lumen and the systemic exclusion of pathogens (Chelakkot et al., 2018). Intestinal epithelial barrier integrity is governed by tight junction proteins, including the Claudin family of proteins (Günzel and Yu, 2013; Lu et al., 2013; Milatz et al., 2010; Patel et al., 2012), and is understood to be compromised in conditions such as inflammatory bowel disease (Clayburgh et al., 2004). Murine studies have shown that the expression of CLDN3, a marker of improved epithelial barrier integrity, is absent at the protein level during gestation and emerges after birth upon completion of morphogenesis and exposure to probiotic bacteria (Patel et al., 2012). However, our data shows that CLDN3 was expressed sporadically in the murine intestine before birth at E17.5 and localized to intervillus domains at P7, which suggests that its emergence may not require exposure to probiotic bacteria. Further, CLDN3 was robustly expressed in the human intestinal epithelium as early as week 18 of gestation, indicating that intestinal CLDN3 expression may not require exposure to microbiota or may be only a partial requirement for establishing barrier integrity.

Fucosyltransferase activity, and in turn fucosylation of brush border and mucinous glycoproteins, has been noted to occur after birth in mice, and is thought to be triggered by exposure to *Bacteriodes thetaiotaomicron* or nutrient-derived polyamines (Biol-N’Garagba et al., 2002; Bry et al., 1996; Capano et al., 1994; Dufour et al., 1988). However, our data for UEA1 reactivity, which shows high affinity for fucosylation events, was visible before birth in both the murine and human intestine. This observation deserves future exploration, as it suggests that significant fucosylation is occurring in the fetal intestine at a point when microbial colonization has not yet occurred (Bry et al., 1996; Perez-Muñoz et al., 2017; Schaedler et al., 1965).

Our dataset confirms the well-established observation that completion of intestinal morphogenesis occurs prenatally in humans and postnatally in mice (Chin et al., 2017; McCracken and Lorenz, 2001). Exposure of the fetal intestine to microbiota at birth is considered a major event that shapes multiple facets of intestinal development and maturation. Our data indicates that certain maturation markers that are considered to be associated with microbial colonization of the neonatal gut in the mouse are expressed prior to birth in humans (Bry et al., 1996; Patel et al., 2012), when the intestinal tract is considered to be sterile (Perez-Muñoz et al., 2017; Schaedler et al., 1965). Although the intestine is thought to be sterile before birth, recent studies have asserted that maternal microbes present in the placenta and amniotic fluid may infiltrate the fetal intestine (Aagaard et al., 2014; Collado et al., 2016; Perez-Muñoz et al., 2017), suggesting that bacteria-mediated maturation may begin *in utero*. The sterility of the prenatal intestine continues to be contested, but if prenatal colonization is confirmed, then it will be worth exploring the impacts that microbes have on intestinal maturation during gestation.

This work provides new insights into intestinal development and maturation during human gestation and should serve as a guide for other researchers studying the intestine, including hPSC-derived tissues, to gauge the relative maturation status of their samples. Our work underscores the difference in the timeline of intestinal development between humans and mice, and reinforces the utility of *ex vivo* models using human cells and tissues, such as intestinal organoids to further study intestinal development and maturation in humans (Forster et al., 2014; Hill et al., 2017; Nadkarni et al., 2017; Senger et al., 2018; Spence et al., 2011).

## Experimental Procedures

### Obtaining tissue sections of human and mouse intestine

Access to fixed tissue sections of human intestine from gestation was gained through Hamilton Health Sciences and St. Joseph’s Healthcare Hamilton in Ontario, Canada. Full approval was obtained from the Hamilton Integrated Research Ethics Board (HiREB) for the use of human tissue for research purposes (project #2399). Archived tissue blocks of fetal autopsies and products of conception were chosen by a staff pathologist using their personal record of completed cases. The gestational age of each sample was determined at the time of diagnosis, based on the patient history and gestational age reference values. Only the tissue blocks containing small bowel were pulled by laboratory staff and identified by the gross description captured on the software Meditech.

Murine intestinal samples were obtained from Dr. Tae-Hee Kim’s lab in the Department of Molecular Genetics at the University of Toronto in Toronto, Ontario. The mouse strain used was C57BL/6. Mice were housed in specific-pathogen-free (SPF) barrier facilities. All experimental work and mouse handling was performed and monitored according to protocols approved by The Centre for Phenogenomics (TCP) Animal Care Committee in Toronto, Ontario (AUP #23-0276H). Mice were sacrificed and the small intestine was harvested accordingly from the chosen embryonic and postnatal ages.

### Immunostaining of tissue sections

Formalin-fixed paraffin-embedded (FFPE) tissue sections were stained according to protocols from R&D Systems for fluorescent IHC staining (https://www.rndsystems.com/resources/protocols/protocol-preparation-and-fluorescent-ihc-staining-paraffin-embedded-tissue) and chromogenic IHC staining (https://www.rndsystems.com/resources/protocols/protocol-preparation-and-chromogenic-ihc-staining-paraffin-embedded-tissue). For chromogenic IHC staining, reagents from Anti-mouse (CTS002), Anti-rabbit (CTS005), and Anti-rat HRP-DAB Cell & Tissue Staining Kit (CTS017) from R&D Systems were used.

Primary antibody information:

| **Antibody/protein** | **Vendor** | **Catalog no.** | **Usage** |
| --- | --- | --- | --- |
| CLDN3 (h/m) | Abcam | ab15102 | 1:100 |
| DEFA5 (h) | Novus Biologicals | NB110-60002SS | 10 µg/ml (1:235) |
| DEFA1 (m) | Cloud-Clone Corp | PAB705Mu01 | 10 µg/ml (1:79) |
| Ki67 (h) | Cell Signaling | 9449 | 1:400 |
| OLFM4 (h) | Abcam | ab85046 | 5 µg/ml (1:120) |
| SI (h/m) | Santa Cruz | sc-27603 | 1:100 |
| UEA1 (h/m) | Vector Laboratories | DL-1067 | 1:100 |

H&E, AB-PAS and chromogenic IHC staining for CDX2 and CHGA were performed by technical staff in the HRLMP at St. Joseph’s Hospital in Hamilton, Ontario, Canada. Antibody information:

CDX2: Clone - DAK-CDX2; Isotype - IgG1, kappa
CHGA: Clone - DAK-A3; Isotype - IgG2b, kappa

### Imaging and processing

IF-stained sections were viewed on an Olympus IX81 inverted fluorescence microscope, and photographs were captured using a Hamamatsu C11440 digital camera. IHC-stained sections were scanned on an Aperio ScanScope slide scanner. Image processing and channel merging was done on ImageJ.

## Author Contributions

R.R.N., J.S.D., and A.A.L. conceived and designed the study, and wrote the manuscript. A.A.L. and R.R.N. performed experimental work and prepared the figures. B.C.C. identified and provided tissue sections of gestational human intestine, and gave conceptual advice. A.A.L., R.R.N., B.C.C., and J.S.D. analyzed the data. J.S.D. supervised the project.

## Acknowledgments

This work was funded by grants from the Canadian Institutes of Health Research (#130499) and the Department of Pathology and Molecular Medicine at McMaster University (#038) awarded to J.S.D. and B.C.C., respectively. Funds provided by the Farncombe Family Digestive Health Research Institute at McMaster University also supported this work. J.S.D. is supported by a Canada Research Chair. We are very grateful to Dr. Tae-Hee Kim and his postdoctoral fellow Dr. Ji-Eun Kim from the Department of Molecular Genetics at the University of Toronto for providing gestational murine intestinal tissue sections. We thank Dr. Michael G. Surette from the Farncombe Family Digestive Health Research Institute at McMaster University for providing important insights.

## Supplemental Information

Supplemental Information includes four figures and their figure legends, and can be found online with this article.

