## Supplemental Information for "Comparison of human and mouse fetal intestinal tissues reveals differential maturation timelines"

#### Supplemental Figure Legends

Figure S1. Chromogenic IHC staining of gestational and postnatal small intestine in humans versus mice for CDX2. Scale bars, 50 µm.

Figure S2. Haematoxylin and eosin (H&E) as well as Alcian blue/periodic acid-Schiff (AB-PAS) staining of gestational and postnatal small intestine in humans versus mice. Scale bars, 50 µm.

Figure S3. Chromogenic IHC staining of gestational and postnatal human small intestine for CHGA. Scale bars, 50 µm.

Figure S4. Fluorescent IHC staining of gestational and postnatal small intestine in humans for Ki67. Scale bars, 50 µm.

#### Supplemental Figures

Supplemental Figures begin on the following page.

### Figure S1

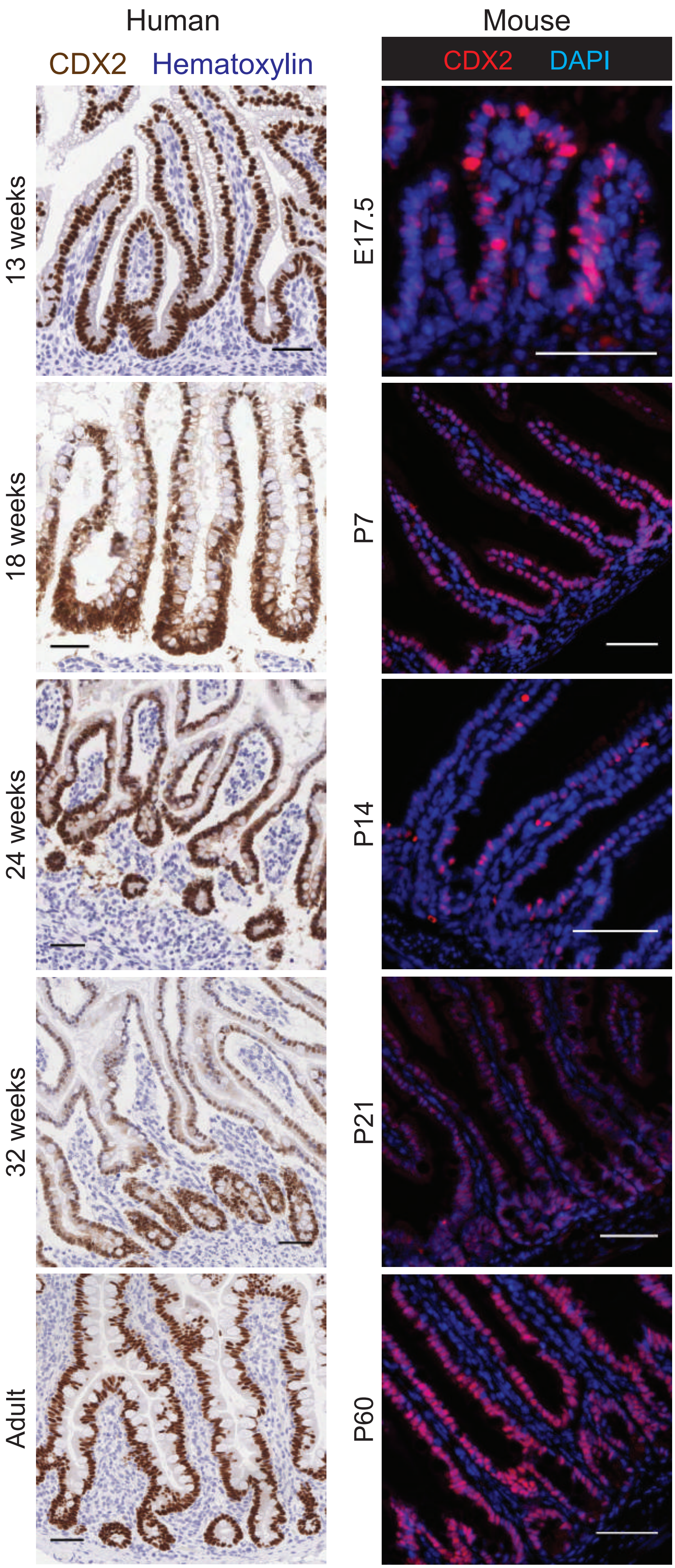

Human
CDX2 Hematoxylin
Mouse
CDX2 DAPI
13 weeks
18 weeks
24 weeks
32 weeks
Adult
E17.5
P7
P14
P21
P60

### Figure S2

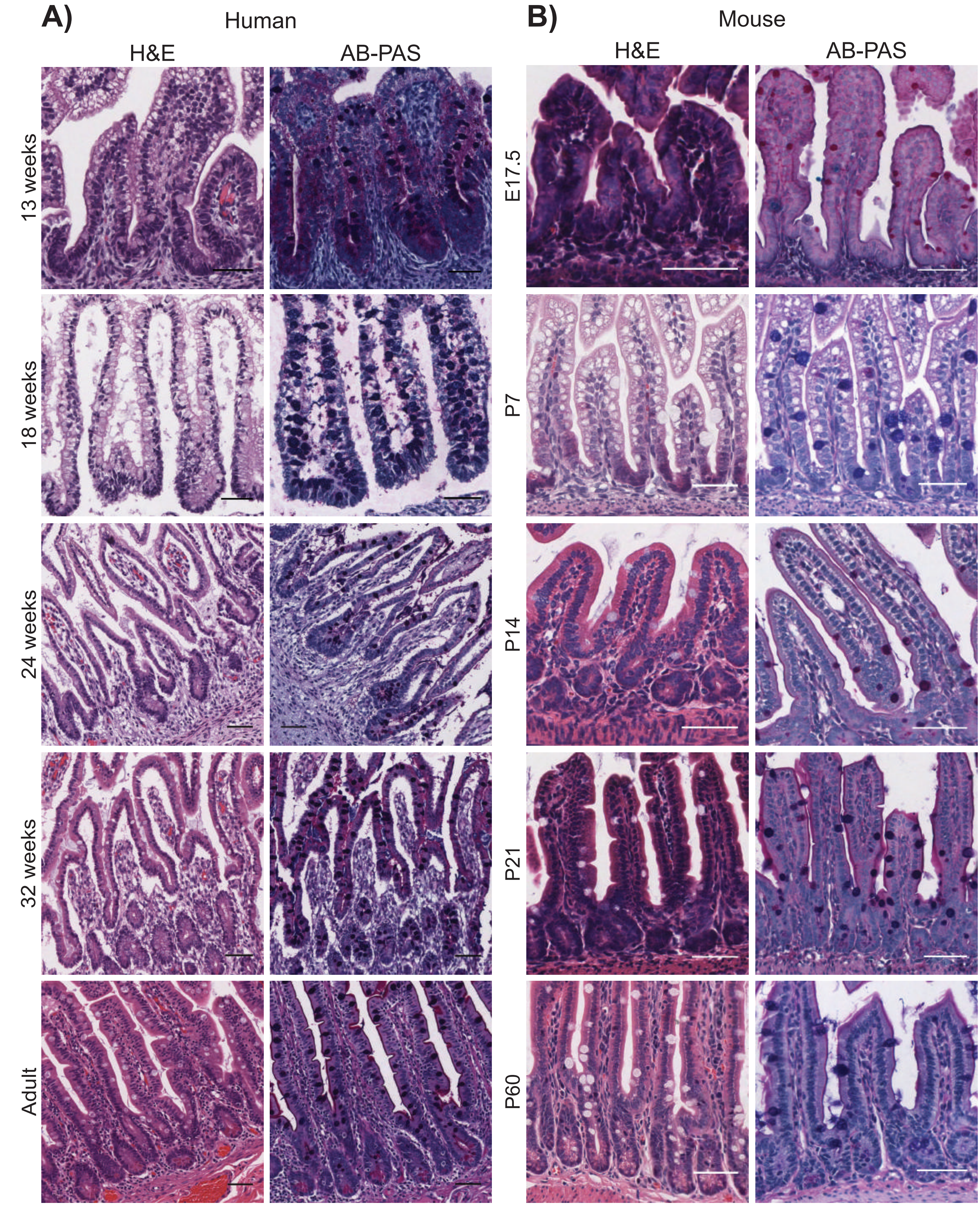
A)
Human
H&E
AB-PAS
13 weeks
18 weeks
24 weeks
32 weeks
Adult
B)
Mouse
H&E
AB-PAS
E17.5
P7
P14
P21
P60

### Figure S3

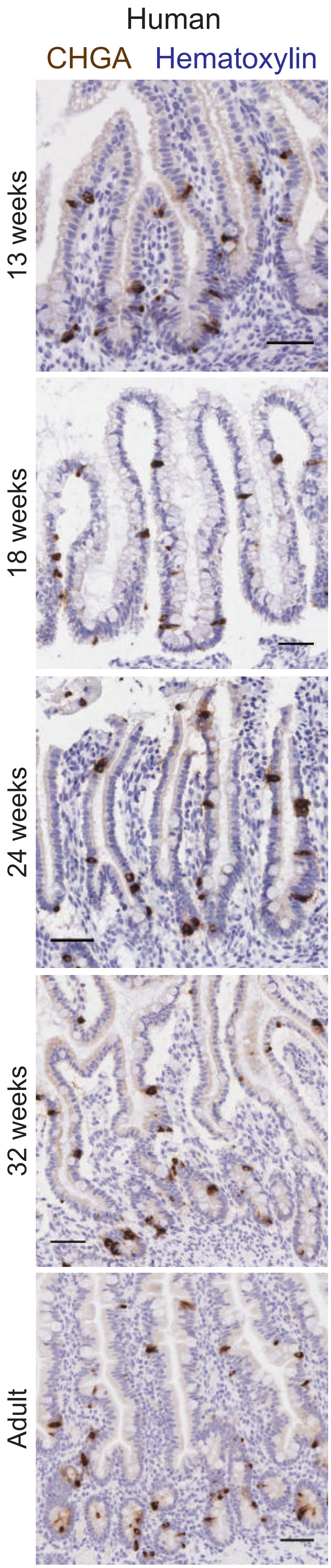
Human
CHGA Hematoxylin
13 weeks
18 weeks
24 weeks
32 weeks
Adult

### Figure S4

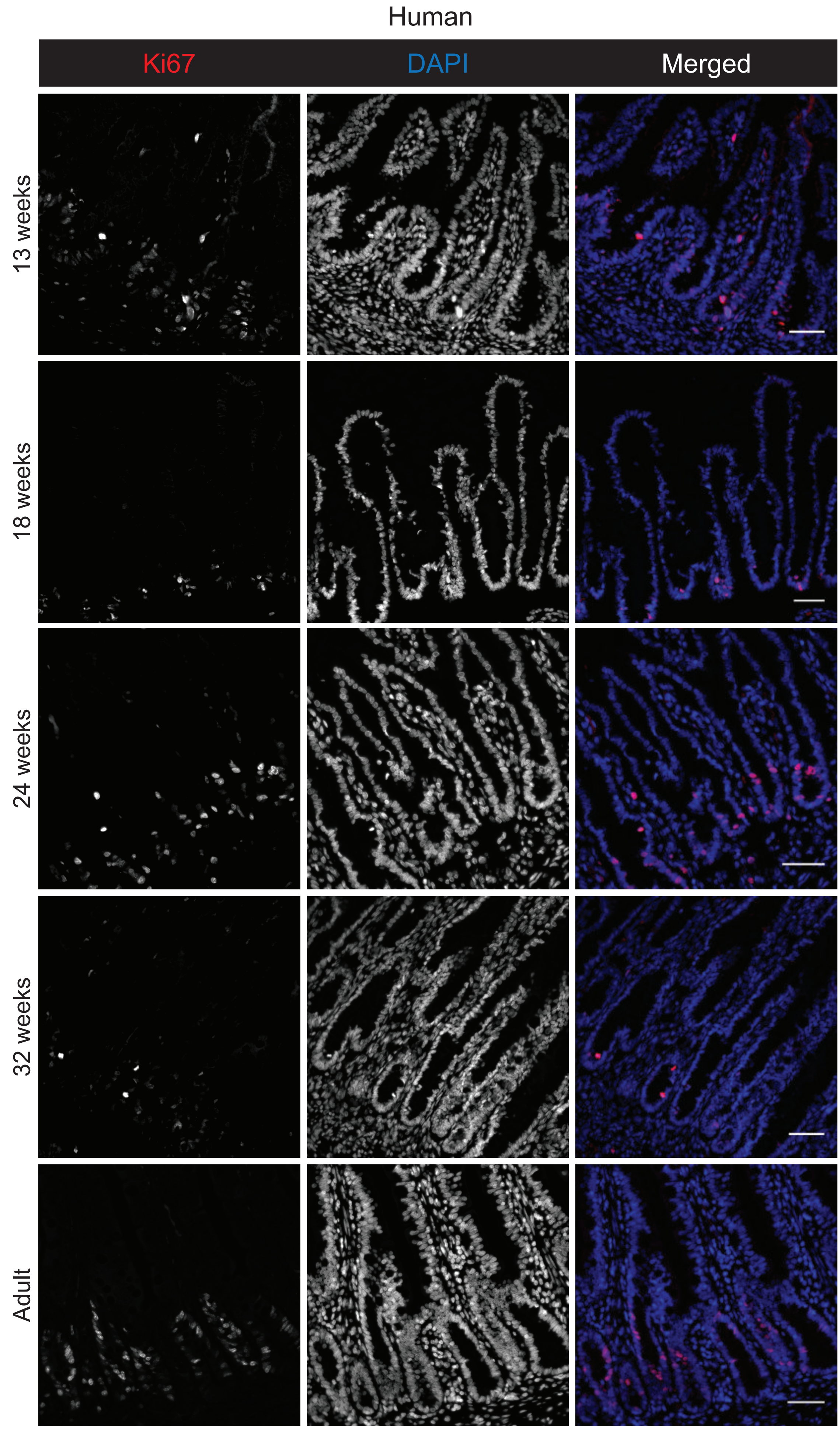

Human
Ki67
DAPI
Merged
13 weeks
18 weeks
24 weeks
32 weeks
Adult
